# Tracking Adoptive T Cell Therapy Using Magnetic Particle Imaging

**DOI:** 10.1101/2020.06.02.128587

**Authors:** Angelie Rivera-Rodriguez, Lan B. Hoang-Minh, Andreina Chiu-Lam, Nicole Sarna, Leyda Marrero-Morales, Duane A. Mitchell, Carlos Rinaldi

## Abstract

Adoptive cellular therapy (ACT) is a potent strategy to boost the immune response against cancer. ACT is an effective treatment for blood cancers, such as leukemias and lymphomas, but faces challenges treating solid tumors and cancers in locations like the brain. A critical step for success of ACT immunotherapy is achieving efficient trafficking of T cells to solid tumors, and the non-invasive and quantitative tracking of adoptively transferred T cell biodistribution would accelerate its development. Here, we demonstrate the use of Magnetic Particle Imaging (MPI) to non-invasively track ACT T cells *in vivo*. Labeling T cells with the superparamagnetic iron oxide nanoparticle tracer ferucarbotran did not affect T cell viability, phenotype, or cytotoxic function *in vitro*. Following ACT, ferucarbotran-labeled T cells were detected and quantified using MPI *ex vivo* and *in vivo*, in a mouse model of invasive brain cancer. Proof-of-principle *in vivo* MPI demonstrated its capacity to detect labeled T cells in lungs and liver after intravenous administration and to monitor T cell localization in the brain after intraventricular administration. *Ex vivo* imaging using MPI and optical imaging suggests accumulation of systemically administered ferucarbotran-labeled T cells in the brain, where MPI signal from ferucarbotran tracers and fluorescently tagged T cells were observed. *Ex vivo* imaging also suggest differential accumulation of nanoparticles and viable T cells in other organs like the spleen and liver. These results support the use of MPI to track adoptively transferred T cells and accelerate the development of ACT treatments for brain tumors and other cancers.

## INTRODUCTION

Harnessing the immune system has immense potential for cancer treatment. Several strategies have been explored to boost the immune response against cancer, including cancer vaccines, oncolytic viruses, immune checkpoint blockade, and adoptive cellular therapy (ACT). The last two strategies have shown unprecedented clinical responses and first-in-kind approvals in recent years.^1^ In fact, adoptive T cell therapy following lymphodepletive conditioning regimens has emerged as one of the most effective treatment strategies against advanced malignant melanoma, with remarkable objective clinical responses in greater than 70% of patients with refractory metastatic disease. This has included a greater than 40% response rate in patients with brain metastases, demonstrating that the central nervous system (CNS) is not refractory to effective treatment with systemically administered T cells.^2^ Importantly, immunologic surrogate endpoints that correlate with treatment outcome have been identified in these patients, with clinical responses being dependent on the localization of transferred T cells to sites of invasive tumor growth and the persistence of tumor-specific effector memory T cells at high precursor frequency in the blood.^3–6^ Despite ACT being successful against melanomas and blood cancers such as leukemia and lymphoma, this therapy has faced challenges in treating solid tumors and invasive CNS malignancies such as malignant gliomas. One contributing factor lies in the relatively poor trafficking and persistence of T cells in solid tumors following systemic delivery. Therefore, technologies that enable the non-invasive and quantitative tracking of adoptively transferred T cells would have tremendous impact in accelerating development of new and effective ACT strategies.

Biomedical imaging can be used to track cell therapies, commonly using luminescence/fluorescence imaging, magnetic resonance imaging (MR), positron emission tomography (PET), or single-photon emission computed tomography (SPECT). Cell tracking using these techniques generally follows two approaches: (i) genetic modification of cells to express a marker that enables optical visualization (IVIS^®^) or nuclear tomographic imaging (PET/SPECT) ^7^ and (ii) labeling of cells using agents that provide contrast (MR)^8^ or generate a signal suitable for imaging (IVIS^®^, PET/SPECT).^9^ However, these current imaging technologies possess several disadvantages, such as low cell detection sensitivity, limited tissue penetration depth, low resolution, and artifacts that preclude the unambiguous tracking of cells.

Magnetic particle imaging (MPI) is a novel imaging technology that enables the non-invasive visualization of the distribution of biocompatible superparamagnetic iron oxide (SPIO) tracers.^10–11^ MPI was first reported in 2005^10^ and is rapidly progressing towards clinical translation.^12^ There is no tissue background with MPI, and signal intensity is proportional to the SPIO mass.^13^ The signal in MPI does not suffer from the artifacts arising from hypointense SPIO signals observed with MR. Further, the signal in MPI is not attenuated by tissue and has no practical tissue penetration limitations. MPI does not use ionizing radiation and SPIO tracers have a long shelf life *ex vivo*, while they break down *in vivo*, with iron being incorporated into hemoglobin and ferritin.^14–15^ In addition, images obtained from MPI can be co-registered with other modalities, like computed-tomography (CT) and MR. Proof-of-principle studies have explored the application of MPI for blood pool imaging,^16–20^ lung perfusion,^21–23^ bleed detection,^24–25^ and tracking of nanoparticle accumulation in cancer tumors.^26–27^. To date, cell-tracking studies using MPI have focused almost exclusively on stem cells. ^11, 28–29^

Here, we evaluate the potential of MPI to track ACT by labeling T cells *ex vivo* with the commercially available MR tracer, ferucarbotran. Ferucarbotran is manufactured by Meito Sangyo (Japan) under the same conditions as Resovist and is marketed in the US as VivoTrax™ by Magnetic Insight for use in MPI. While ferucarbotran is not optimized for MPI, it has been adopted as a reference standard in the MPI literature due to its availability. We assess the effects of ferucarbotran labeling on viability, phenotype, and effector function of tumor-specific T cells. Intracellular localization of ferucarbotran is evaluated using complementary microscopy techniques. T cell uptake of ferucarbotran is quantified using MPI and validated against a well-established iron quantification assay. Linearity between MPI signal and labeled T cell count is demonstrated. *In vivo* MPI shows T cell accumulation in the lungs, followed by redistribution to other organs when administered systemically. *Ex vivo* co-localization of MPI signal and a T cell fluorescence reporter suggests that ferucarbotran labeled T cells accumulate in the brain of tumor bearing mice. For mice receiving T cells intra-ventricularly, *in vivo* MPI permits tracking localization over time. Therefore, our proof-of-principle studies demonstrate that ferucarbotran labeled T cells can be imaged non-invasively using MPI.

## METHODS

### Animals and cell lines

C57BL/6 wild type mice were obtained from Jackson Laboratory. Transgenic pmel specific DsRed C57BL/6 mice were obtained from breeding a DsRed transgenic mouse (B6.Cg-Tg(CAG-DsRed*MST)1Nagy/J) and a pmel transgenic mouse (B6.Cg-Thy1a/Cy Tg(TcraTcrb)8Rest/J) to obtain the DsRed pmel-specific mouse colony. The pmel-specific T cell receptor of these mice recognizes the gp100 antigen expressed by the studied murine glioblastoma cell line, and the DsRed fluorescence allows *ex vivo* identification of transferred T cells. The murine glioblastoma cell line KR158B-Luc was a kind gift from Tyler Jacks.^30^ The KR158B-Luc cell line was genetically modified to express gp100 (Kluc-gp100) using the lentiviral vector pD2107-CMV-gp100. Kluc-gp100 cells were cultured in Dulbecco’s Modified Eagle Media (DMEM) containing 10% fetal bovine serum (FBS). All animals were housed in specific pathogen-free facilities. Experiments were performed according to University of Florida Institutional Animal Care and Use Committee (IACUC) approved protocols.

### T cell isolation and activation

Spleen was harvested from 4-to 8-week-old pmel DsRed transgenic mice. Cells were cultured in Roswell Park Memorial Institute (RPMI) 1640 (Gibco, Waltham, MA) medium supplemented with 10% FBS, 1% non-essential amino acids, 1% L-glutamine, 1% sodium pyruvate, 0.1% 2-mercaptoethanol, 1% penicillin/streptomycin, and recombinant human IL-2 (50 U/mL, R&D Systems, Minneapolis, MN, USA). Single cell suspensions were activated with 1 μg/ml Concavalin A (Sigma, St. Louis, MO) at days 1 and 4 after isolation. ^31^

### Cell Labeling and cell viability

Pmel DsRed T cells were exposed to different concentrations of ferucarbotran nanoparticles (25 μgFe/ml to 200 μgFe/ml) (Meito Sangyo Co., LTD, Japan) at day 7 after isolation for 24 h in T cell media supplemented with IL-2. After exposure, T cells were centrifuged at 500 rcf for 5 minutes at 4 °C, supernatant was discarded and cells were washed 3x with PBS by repeated centrifugation. Cells were resuspended in fresh T cell media. Cell viability was evaluated by two methods. DsRed fluorescence was measured in a plate reader (SpectraMax M5 Microplate Reader, Molecular Devices, LLC, San Jose, CA, USA) at 540 nm Ex/590 nm Em for the different groups of labeled cells. Trypan blue exclusion assay was used to quantify live cells after labeling. Data was normalized relative to control group.

### Flow cytometry

T cells were washed with PBS containing 2% FBS prior to the addition of antibodies. Cells were stained with fluorophore-conjugated antibodies specific for mouse CD3-AF700 (BD Biosciences, San Jose, CA, USA), CD8-APC (eBioscience, Waltham, MA), CD27-PE (BD Biosciences, San Jose, CA, USA), and CD44-BV421 (BD Biosciences, San Jose, CA, USA) in PBS with 2% FBS for 15 min at 4 °C. Cells were then fixed in 4% paraformaldehyde for 10 min at 4 °C and washed twice in PBS with 2% FBS. Flow cytometry was performed on a Canto II flow cytometry system (BD Biosciences, San Jose, CA). ^31^

### Functional assays

To determine if T cells remain functional after labeling with ferucarbotran, pmel DsRed T cells were co-cultured without IL-2 at a 5:1 ratio with KR158B-Luc gp100 glioblastoma cell line for 24 h. Levels of luciferase production from glioblastoma cells were measured using D-luciferin (150 μg/ml final concentration, Perkin Elmer, Waltham, MA, USA) and luminescence measurement in a plate reader after 15 minutes. T cell release of interferon gamma (IFN-γ) was evaluated after labeling. For this experiment, cells were co-cultured at the same conditions stated above in a round-bottom 96-well plate, after 24 h supernatant was collected by centrifuging the plates at 200 rcf for 5 minutes at 4 °C. Mouse IFN-γ Platinum ELISA (Invitrogen, Carlsbad, CA, USA) was performed on the supernatants.^31^

### Microscopy

Labeled and unlabeled T cells were air dried in a microscopy slide, followed by fixation in 4% paraformaldehyde and staining with Prussian Blue (Sigma, St. Louis, MO, USA) to image ferucarbotran deposits within the cell via optical microscopy. Fluorescence microscopy (Keyence Epi-Fluorescence microscope BZ-X710, Keyence, Japan) was performed using anti-dextran FITC antibody (Stemcell Technologies, Vancouver, Canada) co-incubated with T cells for 24 h in a chamber slide for live cell imaging. Cells were stained with Hoechst 33342 for 5 minutes followed by live cell imaging microscopy. To further evaluate ferucarbotran uptake, transmission electron microscopy of labeled T cells was performed. Cells were fixed for 24 h in a mixture of 4% glutaraldehyde and 1% paraformaldehyde. Then the sample was processed through a serial step dehydration process with ethanol, stained with osmium, and embedded in epoxy. Blocks were sectioned into 70 nm thin slices using a Leica Ultracut UCT ultramicrotome (Leica Microsystems, Wetzlar, Germany). Thin sectioned slices were dried in a copper grid prior to imaging in Hitachi H7600 electron microscope (Hitachi High-Technologies Corporation, Japan) and FIB - FEI Helios EDAX EDS/EBSD (Field Electron and Ion Company, Hillsboro, Oregon, USA).

### Iron Quantification

Two methodologies were used to quantify iron uptake: 1,10-phenanthroline assay and MPI. For the 1,10-phenanthroline colorimetric assay, 2×10^6^ cells were digested in nitric acid at a concentration of 5×10^5^ cells/ml at 115 °C until nitric fully evaporated. The sample crust that is formed was dissolved in water and reacted with hydroxylamine for 1 h, followed by addition of sodium acetate and 1,10-phenanthroline. A calibration curve was prepared alongside the samples and absorbance was measured at 508 nm. For MPI iron quantification, 2×10^6^ cells were centrifuged in a 0.2 mL centrifuged tube and the pellet was immobilized with 4% agar solution prior to measurement in the MPI. Three fiducials containing approximately 100%, 50% and 10% of the total MPI signal were used within the same field of view (FOV) on the MPI. Longitudinal isotropic scans were taken using the Momentum™ imager from Magnetic Insight (Alameda, CA, USA). Images were analyzed using VivoQuant™ software, and iron uptake was determined using the known iron mass in the fiducials.

### Brain tumor implantation and ACT

Murine KR158B-Luc-gp100 glioblastoma cells were cultured as adherent monolayers and harvested with 0.05% trypsin. Tumor cells were resuspended in 1x PBS and mixed with methylcellulose (1:1 ratio) (R&D Systems). 8- to 10-week old naïve C57BL/6 mice were anesthetized with isoflurane and placed in a stereotactic frame. Intracranial implantation of 10^4^ KR158B-Luc-gp100 glioblastoma cells in 2.5 μL was performed 2 mm to the right of the bregma and 4 mm below the skull using a 25G needle attached to a 250 μL syringe (Hamilton, Reno, NV). Mice were monitored and sacrificed before reaching endpoint. ^31^ Ferucarbotran-labeled activated pmel DsRed T cells (10×10^6^ per mouse), were washed and resuspended in 100 μL PBS, and then injected intravenously via the tail vein. For intraventricular injections, T cells (1×10^6^ per mouse) were suspended in 5 μL PBS. Mice were anesthetized with isoflurane and placed in a stereotactic frame. 5×10^5^ cells were implanted at ±1 mm mediolateral, −0.3 mm anteroposterior, and −3 mm dorsoventral, in a volume of 2.5 μL in each of the two lateral ventricles. To improve engraftment of T cells, animals were lympho-depleted with total body irradiation using 500 rad (5 Gy) 24 h prior ACT.

### *In vivo* and *ex vivo* imaging

Mice were anesthetized with isoflurane (Patterson Companies, Inc, St. Paul, MN, USA), and 150 mg Luciferin/kg body weight were injected in the intraperitoneal cavity. Animals were placed on the warmed In Vivo Imaging System (IVIS^®^ Spectrum CT, Perkin Elmer, Waltham, MA, USA) stage under 1.5% isoflurane and luminescence acquisition was taken 15 minutes after injection. Epi-fluorescence images using DsRed (570/620 nm) were also obtained. After IVIS^®^, the mouse was transferred to the MPI bed under anesthesia, and images were acquired using high sensitivity mode (3 T/m) in a FOV of 6 × 12 cm. Fiducials of known iron mass were within the same FOV in the MPI. The MPI bed was adapted to fit the CT stage to perform dual 3D imaging and anatomical co-registrations. Anatomic CT (IVIS^®^ Spectrum CT, Perkin Elmer, Waltham, MA, USA) reference images were acquired on anesthetized animals (20 ms exposure time, 440 AI X-Ray filter), and MPI/CT were co-registered in the 3D Slicer open source software.^32^ After *in vivo* imaging, mice were euthanized with isoflurane overdose and the liver, lungs, spleen, kidneys, and brain were collected and imaged via MPI and IVIS^®^ epi-fluorescence.

### Statistical Analysis

One-way ANOVA with Tukey’s multiple comparison test were performed to determine statistical significance for *in vitro* experiments. Statistical significance difference was set for p-values less than 0.05. Graph Pad Prism (La Jolla, CA) and Microsoft Excel (Redmond, WA) were used to conduct all analyses. Error bars represent the standard error of the mean.

## RESULTS AND DISCUSSION

### T cell viability, cytotoxic phenotype, and effector function are unaffected by labeling with ferucarbotran

To examine T cell labeling effects on cell viability, phenotype, and function, cytotoxic pmel-specific DsRed T cells were harvested from the spleen of transgenic mice, activated, expanded *ex vivo,* and labeled with ferucarbotran. These pmel transgenic T cells are specific for an epitope of the melanoma tumor antigen gp100, which is expressed by murine B16 melanomas and GL261 gliomas.^33–34^ The clinically relevant intracranial murine glioma cell line KR158B was modified to express luciferase and gp100 (KR158B-Luc-gp100; Kluc-gp100), while retaining the high-grade glioma features of the parental cell line. The KR158B-Luc-gp100 (Kluc-gp100) cell line forms islands of invasive tumor infiltrate that closely resemble those observed in human gliomas and is refractory to radiation and chemotherapy.^35^ Therefore, we used pmel T cells that specifically recognize the gp100 antigen expressed by this Kluc-gp100 glioblastoma cell line.

Ferucarbotran tracer was used to label murine T cells for *in vivo* tracking following transfer in the immunocompetent murine Kluc-gp100 glioblastoma model. Ferucarbotran is a mixture of Fe_2_O_3_ and γ-Fe_3_O_4_ nanoparticles with core size of ~5 nm, in a carboxydextran matrix with hydrodynamic diameter of 40-60 nm and negative ζ- potential.^36–37^ These particles were developed as an MR contrast agent,^36, 38^ and are not optimized for MPI. However, ferucarbotran is available commercially, has reasonable MPI tracer properties, and serves as a platform for proof-of-principle studies and comparisons across groups.

We evaluated the effects of labeling on the viability and phenotype of DsRed T cells incubated with various concentrations of ferucarbotran (0 to 200 μgFe/ml) for 24 h. Cell viability was determined from DsRed fluorescence measurements and trypan blue exclusion assay (**Figure 1A**) and T cell phenotype was evaluated using flow cytometry (**Figure 1B**). For phenotyping, it was previously demonstrated that activation using Concavalin A favors T cell expansion towards CD8+ with CD44+ expression, corresponding to activated T cells ^31^. Therefore, CD8+/CD44+ markers were used to distinguish between activated and naïve T cells, to determine if differences in phenotype were seen after labeling with ferucarbotran. The expression of CD8+/CD44+ on the unlabeled control (n=4), and ferucarbotran-labeled T cells (n=4) was more than 97% on the CD3+ lymphocyte population. CD27+ expression in CD8+ T cells is lost after repetitive antigenic stimulation. ^39^ The expression of CD27+ in both unlabeled and ferucarbotran-labeled T cells is less than 0.2%, suggesting that T cells are effector and not memory T cells. These results show that ferucarbotran labeling does not affect viability or phenotype of pmel DsRed T cells at the studied iron concentrations.

**Figure 1.**
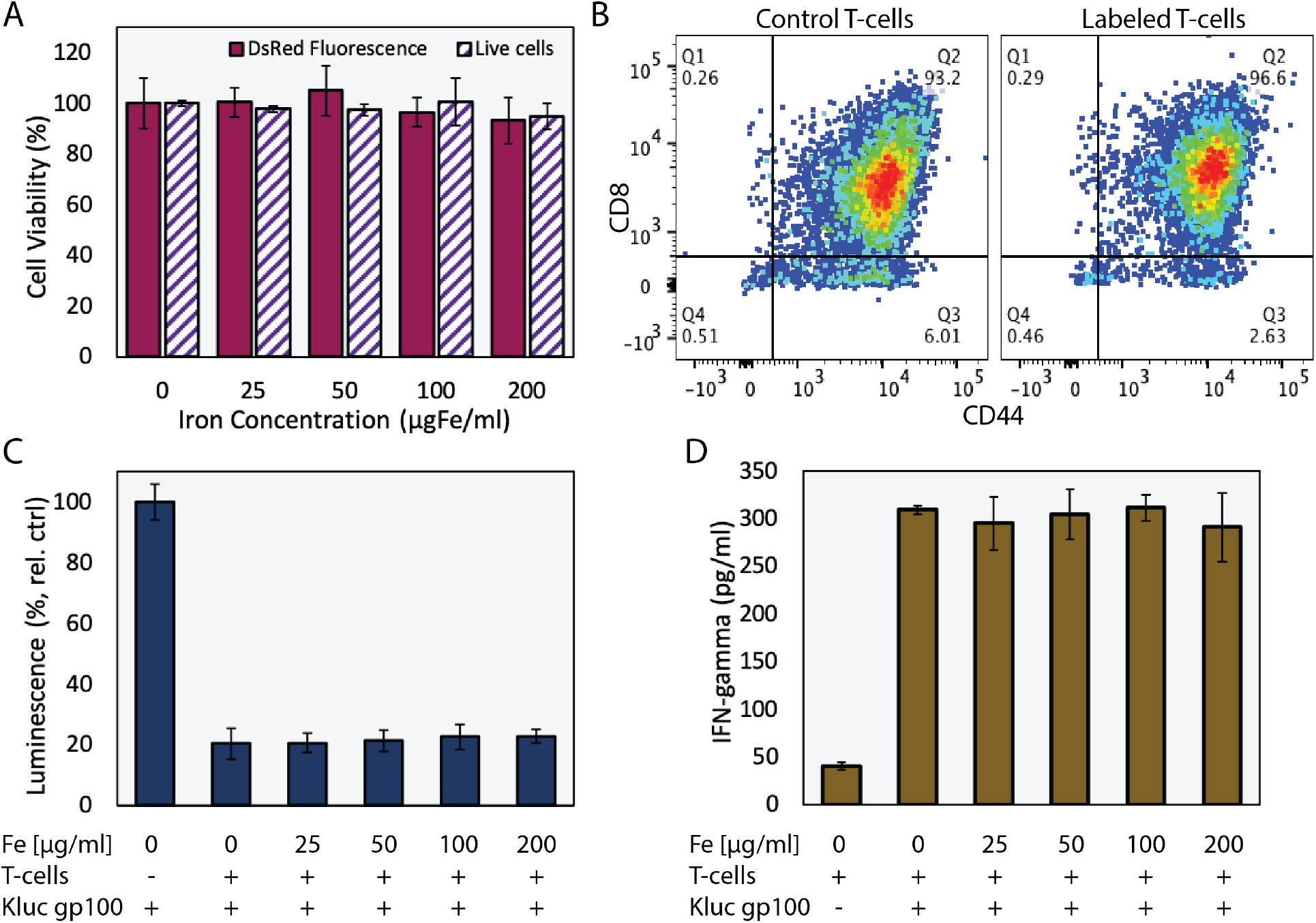
Ferucarbotran labeling does not impair T cells. A) Ferucarbotran-labeled T cell viability relative to control, as determined by DsRed fluorescence measurements and trypan blue exclusion assay. B) Phenotyping of control and labeled (100 ugFe/ml) T cells by flow cytometry. C) Relative viability of Kluc-gp100 cells after co-culture with ferucarbotran labeled T cells, as determined from Kluc-derived luminescence relative to control (no T cells in co-culture). D) IFN-γ release of T cells in co-culture with KLuc-gp100 tumor cells, as determined by ELISA.

T cell effector function was evaluated by quantifying the release of the immunostimulating cytokine interferon-gamma (IFN-γ), which is involved in the anti-tumor cytotoxic T cell response ^31^, and by evaluating cancer cell killing by T cells in co-culture. Cytotoxic activity of ferucarbotran-labeled pmel DsRed T cells against Kluc-gp100 glioblastoma cells was assessed by co-culture of T cells with tumor cells (5:1 ratio) for 24 h. Kluc-gp100 luciferase expression or IFN-γ were measured in separate experiments. A reduction in luciferase expression by the Kluc-gp100 cells in co-culture with T cells would result from a reduction in tumor cell viability due to the cytotoxic activity of the T cells. Results show that labeling T cells using ferucarbotran does not impair their ability to kill Kluc-gp100 cells (**Figure 1C**), or their ability to produce IFN-γ (**Figure 1D**), for all the ferucarbotran labeling conditions used. No statistically significant differences were observed at any given iron concentration in co-cultured groups using one-way ANOVA (p-value less than 0.05).

### Ferucarbotran associates with the T cell membrane and is internalized, leading to measurable MPI signal

The association and uptake of ferucarbotran nanoparticles by T cells was evaluated using complementary microscopy techniques. Prussian blue was used to detect iron deposits in cells using optical microscopy. **Figure 2A** shows iron deposits detected by Prussian blue staining in the ferucarbotran labeled T cells, whereas staining was absent in control cells. Uptake of ferucarbotran was further evaluated using fluorescence microscopy and an anti-dextran FITC antibody specific for the carboxydextran coating of the nanoparticles (**Figure 2B**). Particles within T cells are shown as green in florescence microscopy. Since the pmel T cells endogenously express reporter DsRed (red) fluorescence, live cell imaging was performed and nuclei was stained using Hoechst 33342 (blue). Immunostaining for dextran was observed in the ferucarbotran-labeled group only. Together, Prussian blue and fluorescence microscopy demonstrate labeling of T cells with ferucarbotran.

**Figure 2.**
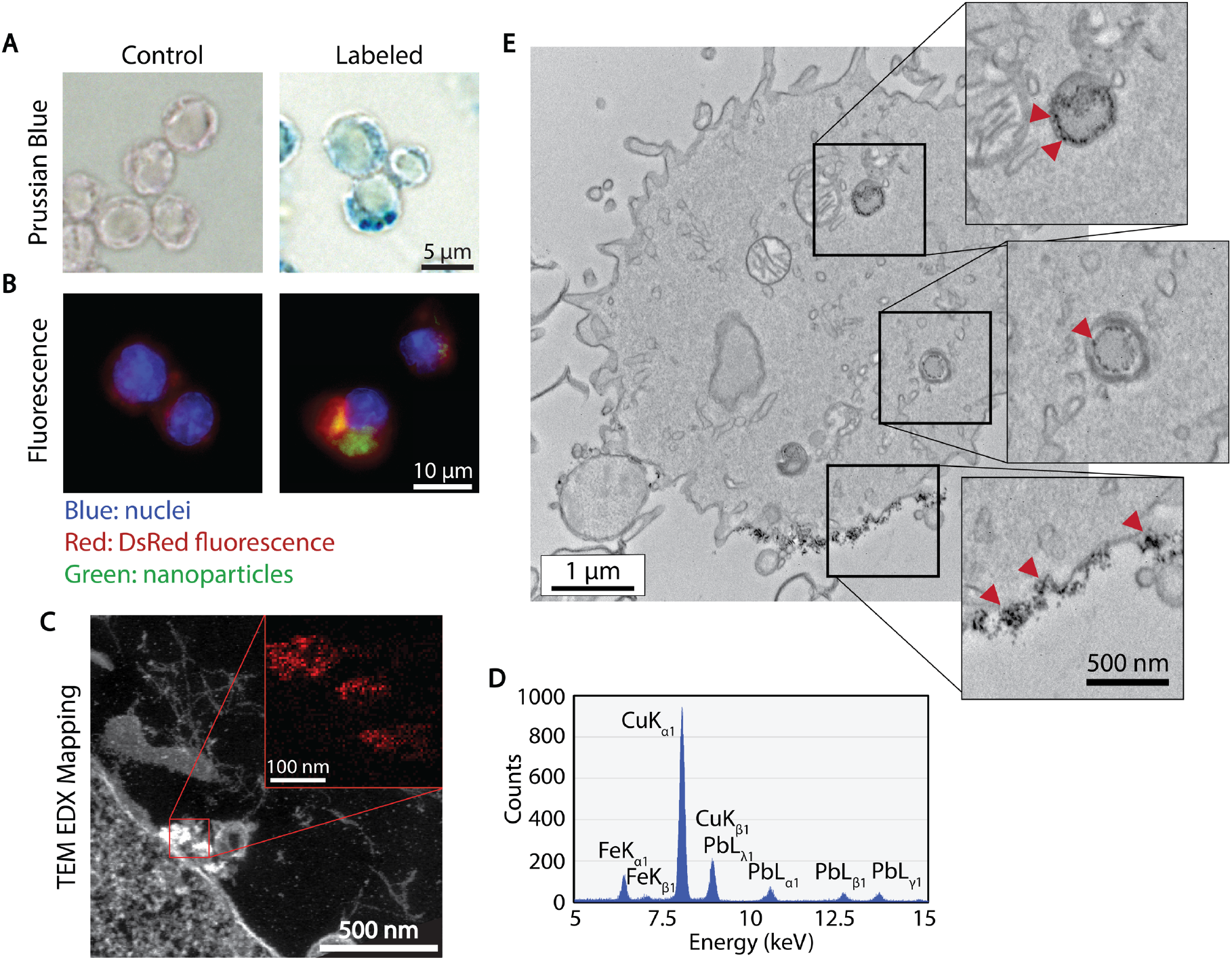
Cellular localization of ferucarbotran tracers in T cells. A) Prussian blue stain for iron oxide. B) Immunofluorescent labeling of ferucarbotran dextran coating (blue-nuclei, red-DsRed fluorescence, green-dextran). C) Z-contrast and energy dispersive X-ray (EDX) spectroscopy show ferucarbotran tracers (bright signal and red in inset) in T cells. D) EDX spectrum shows Fe from ferucarbotran, Cu from grid, and Pb from stain. E) Ferucarbotran tracers are seen in intracellular vesicles (top two insets) and associated with the cell membrane (bottom inset). Cell section stained with osmium only (no Pb) to improve ferucarbotran contrast.

To further evaluate the uptake and intracellular localization of ferucarbotran within the T cells, transmission electron microscopy (TEM) and energy dispersive x-ray (EDX) spectroscopy were used. **Figure 2C** shows z-contrast TEM and elemental mapping of Fe distribution (inset, Fe in red). The EDX spectrum (**Figure 2D**) confirms the presence of iron, copper from the TEM grid, and lead staining for enhanced cell organelles contrast. Since the use of lead staining makes visualization of ferucarbotran inside T cells challenging, ferucarbotran internalization was also confirmed in sections stained only with osmium, to improve contrast between ferucarbotran and cellular components (**Figure 2E**). Ferucarbotran is observed associated with the cell membrane and also in intracellular vesicle-like structures. These results suggest that ferucarbotran nanoparticle might first bind to the T cell membrane, followed by internalization.

### MPI signal quantifies ferucarbotran T cell labeling and is linear with the number of labeled T cells

The extent of T cell labeling with ferucarbotran was quantified using a colorimetric assay and MPI. The 1,10-phenanthroline colorimetric assay, used to quantify iron, involves sample digestion in nitric acid in a two-day process. The working principle of this assay is to form iron (II)- ortho-phenanthroline complexes following iron (III) reduction. Absorbance is then measured at 508 nm. In contrast, ferucarbotran mass can be quantified using MPI in live or dead cells, in a process that takes ~10 min for sample preparation, image acquisition, and analysis. Ferucarbotran uptake was determined using MPI by comparison of the cell sample MPI signal with the signal obtained from a fiducial with free ferucarbotran of known iron mass. We found that ferucarbotran uptake ranges from ~0.4 to ~1 pgFe/cell, depending on the ferucarbotran concentration used during the 24 h incubation period. Of significant interest, the excellent agreement between ferucarbotran uptake determined using MPI or the 1,10-phenanthroline assay suggests that MPI signal per iron mass is not affected after T cell uptake (**Figure 3A**). This result is in contrast to a recent report by Suzuka *et al.* suggesting that ferucarbotran MPI signal is affected after uptake into macrophages or colon cancer cells.^40^ We note that the discrepancy between our observations and those of Suzuka *et al*. may be due to differences in excitation frequency between their prototype scanner (0.4 kHz) and the Momentum™ scanner used here (45 kHz). Ferucarbotran tracer MPI response may be more susceptible to cell uptake at the lower excitation frequencies used in the study by Suzuka *et al.*, in comparison to the higher excitation frequencies used in the present study and previous cell-tracking studies.^28–29^ Additionally, there might be differences in uptake and cellular localization patterns between the T cells used here and the macrophages or colon cancer cells used by Suzuka *et al*.

**Figure 3.**
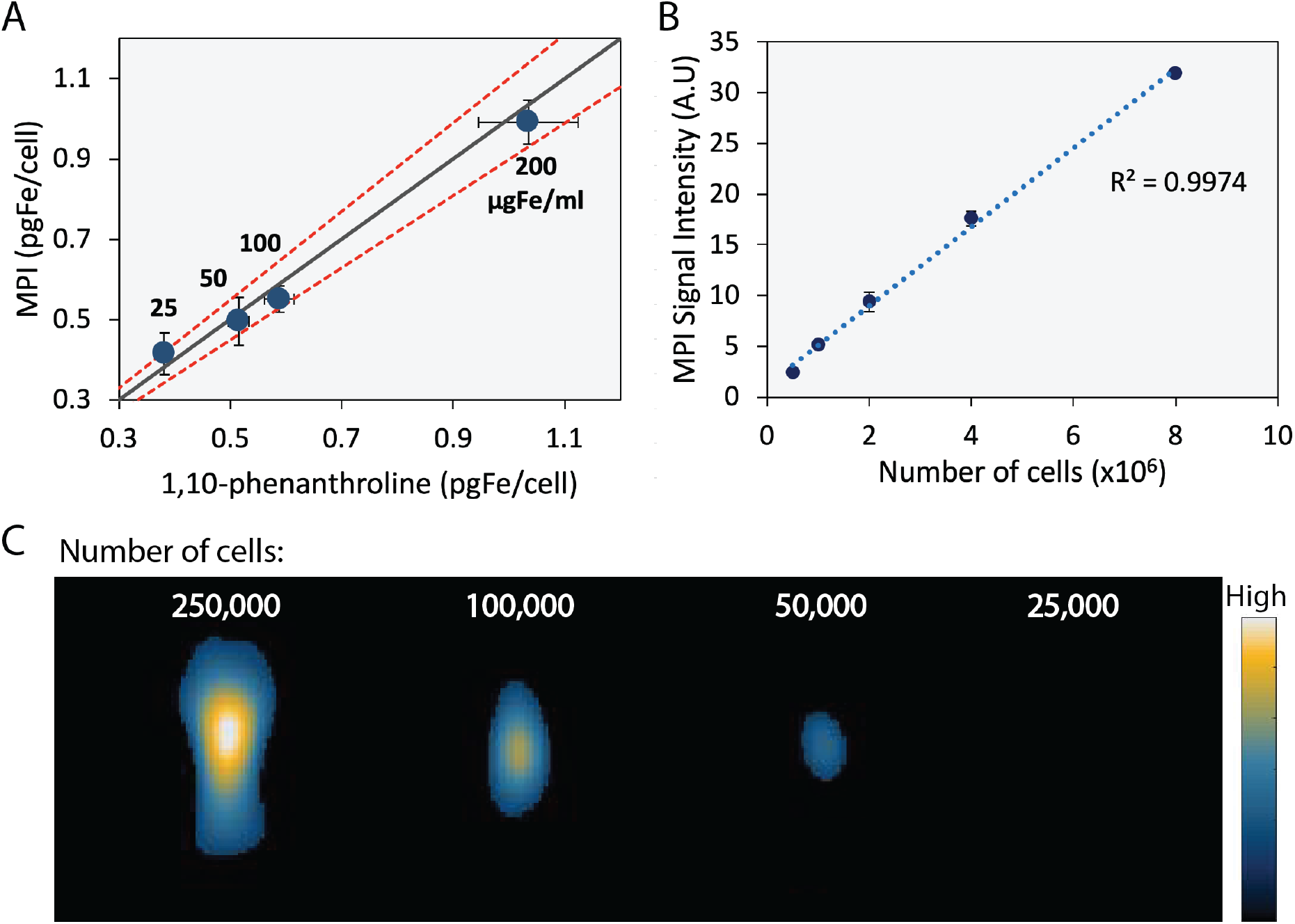
Ferucarbotran-labeled T cells can be quantified using MPI. A) Iron content per cell, quantified using 1,10-phenanthroline assay and MPI. Solid line is parity. Dashed line corresponds to 90% confidence interval. B) Linear correlation between MPI signal intensity and number of ferucarbotran-labeled T cells. Error bars correspond to standard deviation (n=3). Some error bars are smaller than the markers. C) 2D maximum intensity projection scans for samples containing decreasing numbers of ferucarbotran-labeled T cells.

Linearity between MPI signal and number of ferucarbotran-labeled T cells was evaluated. Ferucarbotran mass in the sample was quantified through image analysis using a fiducial with known mass. Excellent linear correlation between MPI signal intensity and number of ferucarbotran-labeled T cells was observed (**Figure 3B**). 2D-MPI scans of samples with different numbers of ferucarbotran-labeled T cells demonstrated that 5×10^4^ cells could be detected in the MPI with only one average per projection (**Figure 3C**). Of note, lower numbers could also be detected, but not quantified, using maximum intensity projection and sample co-registration with optical images. The limit of detection determined for ferucarbotran in our MPI system is approximately 38 ngFe (see Supplementary Information). If each T cell is labeled with 1 pgFe/cell, we should be able to detect approximately 3.8×10^4^ cells in a voxel.

We note that cell detection sensitivity in MPI is determined by the intrinsic signal per iron mass of the tracer used and the extent of cell labeling with the tracer. In the present study the cell detection sensitivity was approximately 50,000 cells. Much lower cell detection sensitivity of approximately 200 cells has been reported for stem cells using MPI.^29^ However, in that report, the stem cells had taken up ~27 pgFe/cell. In contrast, T cell uptake of ferucarbotran was relatively low at ~1 pg/cell. Furthermore, that study used a prototype MPI scanner, rather than the commercially available MPI scanner used here. As such, we expect that better T cell sensitivity can be obtained by increasing T cell labeling with the MPI tracer. Additionally, better cell detection sensitivity can be obtained through use of tracers with higher signal per iron mass. For example, the tracer LS-017, which has been optimized for MPI, is reported to have 6 times higher signal intensity than ferucarbotran ^41^, with further improvements in MPI tracer signal being possible in this relatively new field.

### MPI detects and tracks systemically administered ferucarbotran-labeled T cells *in vivo*

*In vivo* experiments were performed in C57BL/6 mice bearing intracranial KLuc-gp100 tumors. Twelve days after cancer cell implantation, ACT was performed using ferucarbotran-labeled T cells (10^7^ cells in 100 μL) delivered via the tail vein. After injection, T cell biodistribution was monitored with IVIS^®^ using the T cell endogenous DsRed reporter fluorescence, and MPI to detect the ferucarbotran-labeled T cells (**Figure 4**). Using MPI, T cell accumulation was initially detected in the lungs, followed by redistribution after 72 h to other major organs, such as the liver and spleen. In contrast, *in vivo* fluorescence imaging failed to detect live T cells in those organs, due to signal attenuation by tissue/hair using epi-fluorescence. Of note, the IVIS^®^ instrument can also perform 3D fluorescence imaging tomography (a.k.a. transillumination), in which the excitation light source is placed opposite to the camera and closer to the animal for better tissue penetration. This technique could allow the detection of signals at a greater depth. However, transillumination in IVIS^®^ relies on the contrast of the animal with the instrument stage to determine the boundaries of the animal and allow selection of the field of view. The mouse strain C57BL/6, used in the present study has black hair, which prevents the IVIS^®^ software from detecting a mouse on a black stage, limiting the transillumination acquisition. Mouse hair, whether, black, white, gray or brown, absorbs and scatters light in both the visible and near-infrared spectrum. Therefore, when performing optical imaging, it is recommended to shave and/or chemically depilate hair in order to minimize interference with imaging signals. As such, our results illustrate the potential advantages of using MPI to track T cells *in vivo* in whole animals without the need of hair removal and with quantitative capabilities.

**Figure 4.**
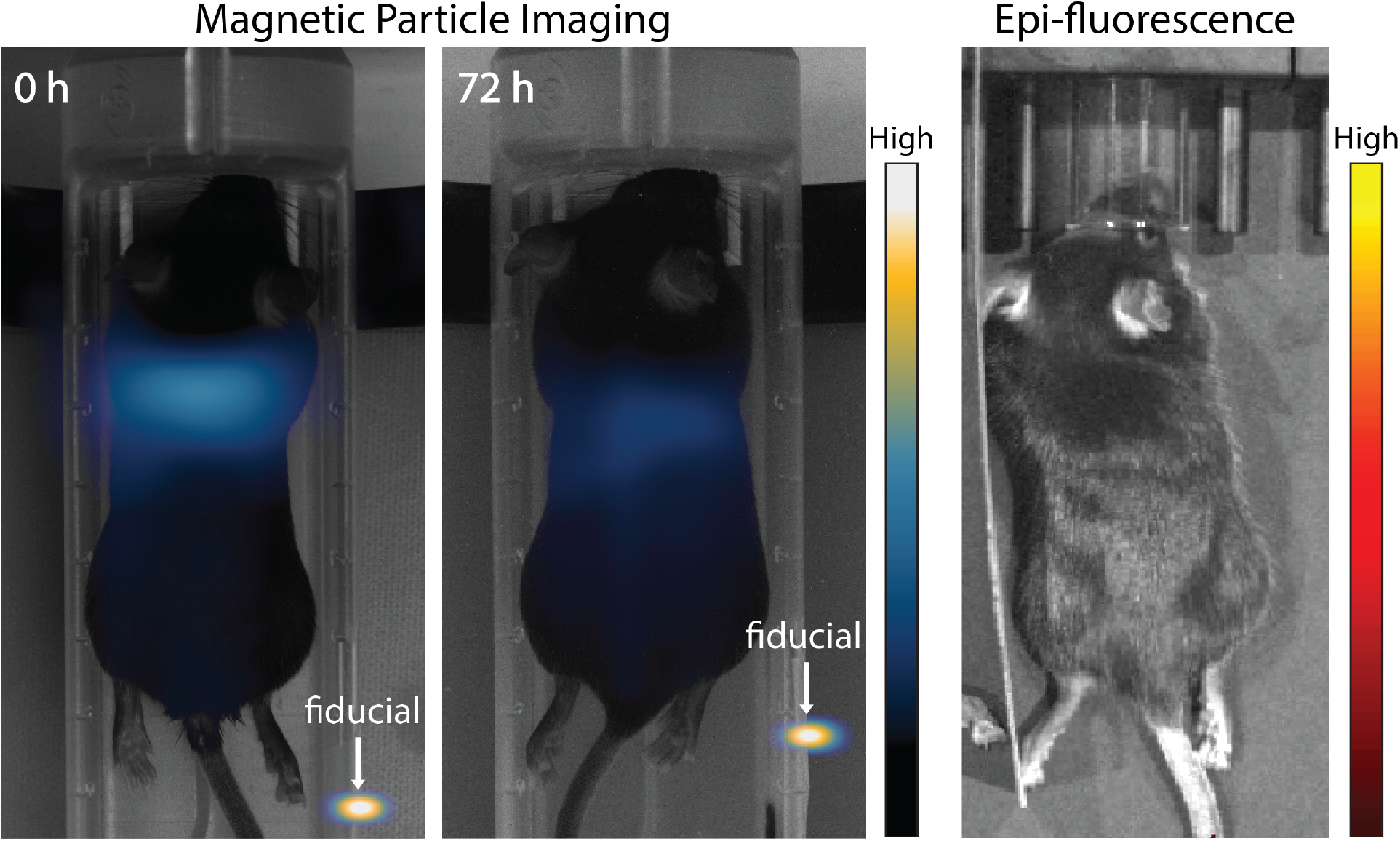
MPI allows for whole-body visualization of ferucarbotran-labeled T cells that are not visible through *in vivo* fluorescence imaging. (Left) MPI taken at 0 h and 72 h after systemic delivery of ferucarbotran-labeled pmel DsRed T cells shows initial accumulation in the lungs followed by redistribution. (Right) Fluorescent imaging of DsRed distribution at 72 h does not show fluorescence due to signal attenuation by tissue/hair for cells distributed in the body.

### *Ex vivo* MPI and fluorescence imaging suggest viable ferucarbotran labeled T cells reach brain tumors following systemic injection

In another study, organs were harvested 24 h after intravenous injection of labeled T cells, and biodistribution was evaluated *ex vivo* using MPI and IVIS^®^. By harvesting the tissue and performing the imaging *ex vivo*, we eliminated disadvantages related to tissue attenuation of optical signal with IVIS^®^. MPI showed accumulation of ferucarbotran-labeled T cells in the liver, spleen, lungs and brain tumor, whereas IVIS^®^ imaging of DsRed fluorescence also detects T cells in the brain tumor and liver (**Figure 5)**. Absence of signal from the lungs and spleen with the IVIS^®^ could be due to cell death after tissue harvesting or low signal sensitivity. Studies using other biomedical imaging modalities to track adoptively transferred T cells have also suggested that T cells accumulate in the liver, lungs, and spleen after intravenous injection.^42–44^ Importantly, observation of both an MPI signal and DsRed fluorescence in the brain tumor suggests that viable ferucarbotran-labeled T cells were able to reach the brain glioblastoma tumor.

**Figure 5.**
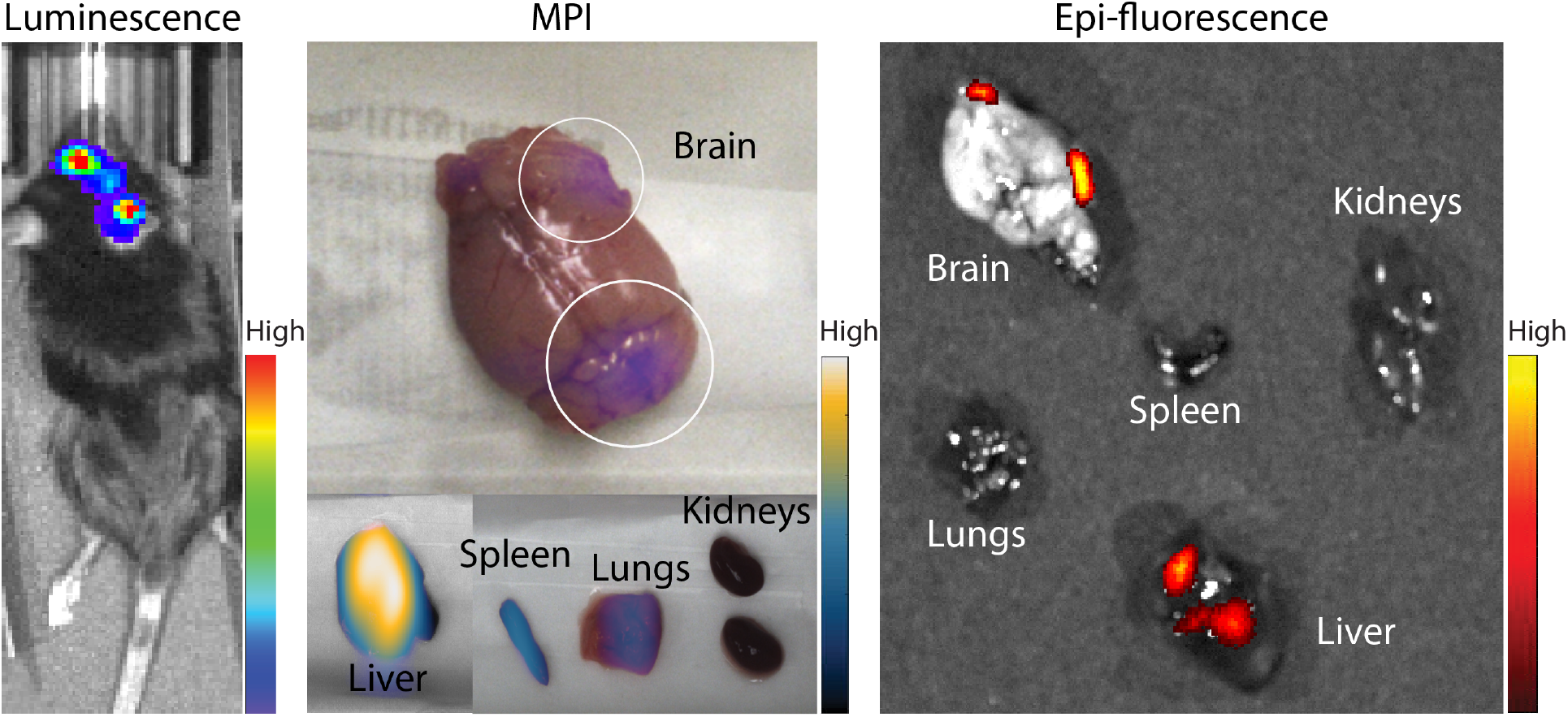
*Ex vivo* MPI and fluorescence imaging suggest that viable ferucarbotran-labeled T cells reach brain tumors 24 h post tail vein injection. *In vivo* luminescence shows the presence of the glioblastoma kluc-gp100 tumor in the brain. MPI shows accumulation of ferucarbotran-labeled T cells in the liver, spleen, lungs, and in two different areas in the brain. Fluorescence imaging shows accumulation of T cells in two areas of the brain, as well as accumulation in the liver. Note that the observation of MPI and DsRed fluorescence signal from T cells in two sites in the brain *ex vivo* is consistent with *in vivo* luminescence imaging that suggests the presence of two tumor masses in those same brain areas.

### Multimodal MPI/CT imaging allows the non-invasive evaluation of T cell distribution in the brain after intraventricular administration

Co-registration of MPI and computed tomography (CT) enabled evaluation of the intracranial localization of ferucarbotran labeled T cells. Intraventricular injections of a known number of T cells were performed in tumor-bearing mice. *In vivo* luminescence showed the presence of Kluc-gp100 brain tumor **(Figure 6)**. T cells (10^6^ cells) were injected in the lateral ventricles, and epi-fluorescence and MPI imaging were performed at different time points after injection. *In vivo* epi-fluorescence signal was not detected at any time point due to signal attenuation by tissue and hair (**Figure 6** shows epi-fluorescence at 36 h). However, T cell DsRed fluorescence signal was detected *ex vivo* after brain harvesting. MPI signal was monitored *in vivo* at different time points, showing only a slight signal decrease over time (Supplementary Information). The change in MPI signal over time was not statistically significant. *In vivo* 3D MPI scans were acquired, followed by CT scans. Anatomical planes demonstrate location and orientation of the MPI signal/labeled T cells in the brain. In addition, 3D anatomical references were obtained from CT imaging, as seen in **Figure 6**. *Ex vivo* MPI was performed, and the MPI signal was co-localized with the *ex vivo* fluorescence signal, suggesting that both signals arise from T cells in the brain.

**Figure 6.**
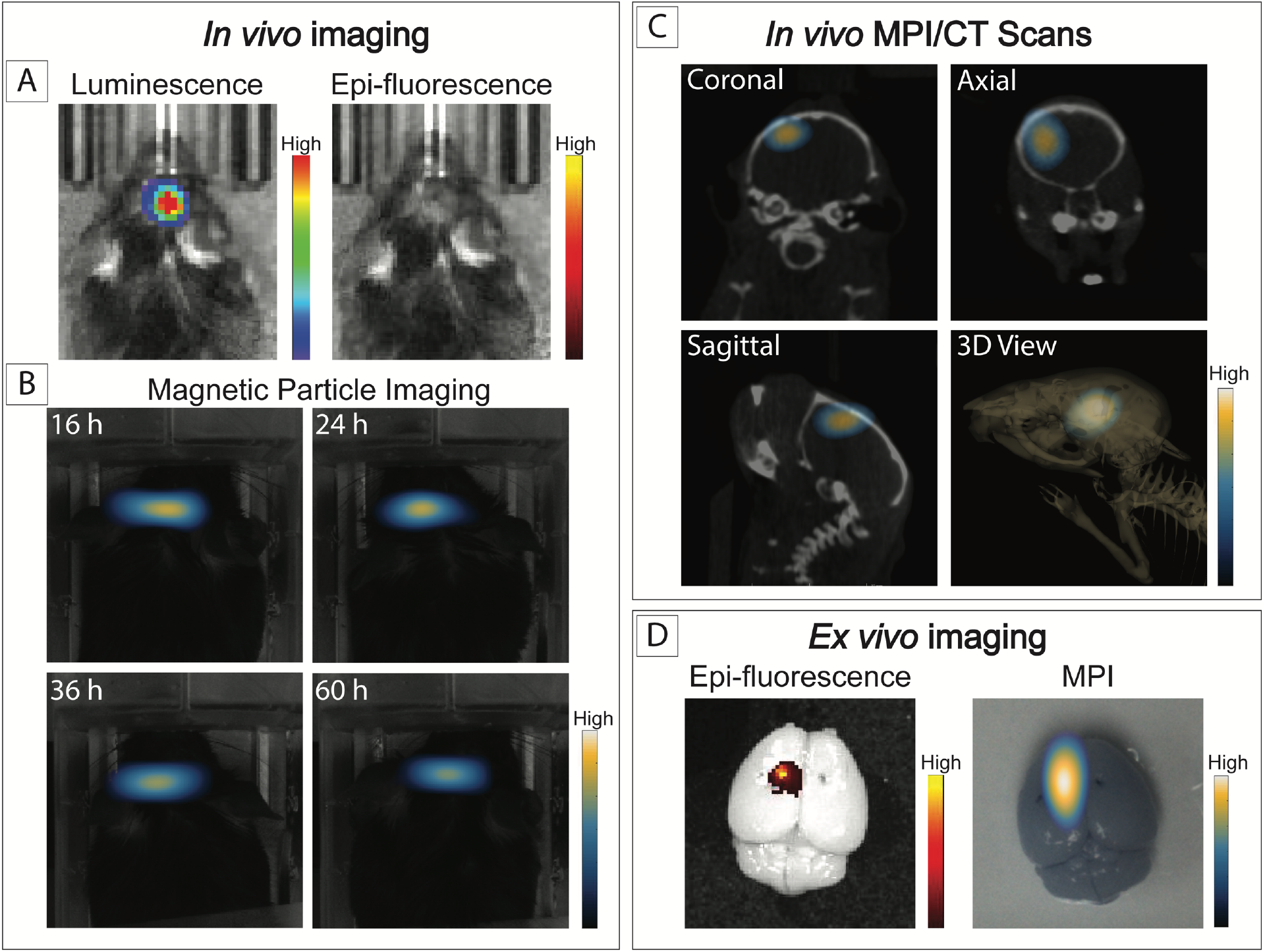
*In vivo* and *ex vivo* imaging after intraventricular administration of ferucarbotran-labeled T cells. A) *In vivo* imaging shows the presence of an intracranial glioblastoma kluc-gp100 tumor (luminescence). After intraventricular injection of labeled T cells, *in vivo* DsRed fluorescence signal is not visible (36 h) due to tissue attenuation. B) Time series of *in vivo* MPI shows a consistent MPI signal at the site of injection. C) MPI co-registered with CT scans, with different anatomical plane cross-sections showing the intracranial localization of ferucarbotran-labeled T cells. D) *Ex vivo* fluorescence signal colocalized with MPI signal.

## CONCLUSIONS

We have demonstrated the use of MPI to non-invasively track the distribution of adoptively transferred T cells labeled with the commercially available superparamagnetic iron oxide, ferucarbotran. Compared with available *in vivo* imaging techniques, MPI provides a unique combination of high sensitivity, resolution, and longitudinal monitoring, with quantification enabled by the linear relationship between signal and tracer mass. These characteristics may help accelerate development and optimization of immune cell therapies by enabling the sensitive and longitudinal tracking of ACT for treatment of refractory malignancies.

In this study, we described the SPIO-labeling of tumor-specific cytotoxic T cells for *in vivo* tracking of adoptive cell therapies. *In vitro* characterizations were performed to determine the extent of ferucarbotran labeling and its effect on cell viability, phenotype and effector function. Ferucarbotran-labeled T cells were not impaired by the labeling process at the studied conditions. MPI permitted the *in vivo* visualization of the biodistribution of ferucarbotran-labeled T cells after systemic and local administration in mice bearing brain tumors. When T cells were administered systemically, MPI noninvasively confirmed entrapment of T cells in the lungs immediately after injection, followed by redistribution to other organs, such as the liver and spleen. Importantly, MPI enabled the non-invasive tracking of T cells after local administration via intraventricular injection, and co-registration of MPI with CT permitted the evaluation of their anatomical localization.

As with other cell tracking techniques that involve the direct labeling of cells, two main challenges of this approach are the dilution of the tracer among the cell progeny and the release of nanoparticles after apoptosis. Furthermore, cell tracking using MPI cannot at present distinguish between live and dead cells, as dead cells might still retain nanoparticles. However, this limitation can be overcome by combining cell labeling using MPI tracers with genetically engineered T cells containing reporter genes to enable multi-modal imaging of their distribution (using MPI) and viability (using luminescence/fluorescence), as shown here. Importantly, our results suggest that viable adoptively transferred T cells reach the brain 24 hours after systemic administration, as shown with *ex vivo* fluorescence imaging for the DsRed reporter gene on the pmel T cells.

To our knowledge, this study is the first to demonstrate the successful labeling of T cells with superparamagnetic iron oxide nanoparticles for use with MPI. Despite the low uptake of nanoparticles by T cells observed in our study (~1 pgFe/cell), we were able to demostrate the non-invasive monitoring of these cells after systemic administration and local delivery. Our results suggest that MPI is an attractive approach to track T cells and provide unique insight into the fate of these cells following adoptive cell therapy for cancer.

## ACKNOWLEDGEMENTS

We thank Dr. Tyler Jacks from Massachusetts Institute of Technology for providing the KR158B-Luc cell line. We thank Dr. Fernanda Pohl-Guimaraes from the UF Brain Tumor Immunotherapy Program in the University of Florida for her input in essential aspects of this study. This work was supported by the National Science Foundation Graduate Research Fellowship Grant No. DGE-1315138 (A.R.R.) and DGE-1842473 (A.R.R.), and NIH/NCI awards 1R01CA175517 (D.A.M.), R01CA195563 (D.A.M.). Research in this publication was supported by the University of Florida Clinical and Translational Science Institute, which is supported in part by the NIH National Center for Advancing Translational Sciences under award number UL1TR001427. The content is solely the responsibility of the authors and does not necessarily represent the official views of the National Institutes of Health.

## CONFLICT OF INTEREST

The authors declare the following competing financial interest(s): D.A.M. has patented immunotherapy related technologies that have been licensed by Annias Immunotherapeutics, Inc., Immunomic Therapeutics, Inc., Celldex Therapeutics, Inc., and iOncologi, Inc. D.A.M. receives research funding from Immunomic Therapeutics, serves as a consultant/advisor to Tocagen, Inc., and is co-founder of iOncologi, Inc., a biotechnology company specializing in immuno-oncology. The other authors declare no conflicts.

## REFERENCES

1. Kelly, P. N., The Cancer Immunotherapy Revolution. Science 2018, 359 (6382), 1344–1345.

2. Phan, G. Q.; Rosenberg, S. A., Adoptive cell transfer for patients with metastatic melanoma: the potential and promise of cancer immunotherapy. Cancer Control 2013, 20 (4), 289–97.

3. Rosenberg, S. A.; Restifo, N. P.; Yang, J. C.; Morgan, R. A.; Dudley, M. E., Adoptive cell transfer: a clinical path to effective cancer immunotherapy. Nat Rev Cancer 2008, 8 (4), 299–308.

4. Zhou, J.; Shen, X.; Huang, J.; Hodes, R. J.; Rosenberg, S. A.; Robbins, P. F., Telomere length of transferred lymphocytes correlates with in vivo persistence and tumor regression in melanoma patients receiving cell transfer therapy. J Immunol 2005, 175 (10), 7046–52.

5. Robbins, P. F.; Dudley, M. E.; Wunderlich, J.; El-Gamil, M.; Li, Y. F.; Zhou, J.; Huang, J.; Powell, D. J., Jr.; Rosenberg, S. A., Cutting edge: persistence of transferred lymphocyte clonotypes correlates with cancer regression in patients receiving cell transfer therapy. J Immunol 2004, 173 (12), 7125–30.

6. Powell, D. J., Jr.; Dudley, M. E.; Robbins, P. F.; Rosenberg, S. A., Transition of late-stage effector T cells to CD27+ CD28+ tumor-reactive effector memory T cells in humans after adoptive cell transfer therapy. Blood 2005, 105 (1), 241–50.

7. Acton, P. D.; Zhou, R., Imaging reporter genes for cell tracking with PET and SPECT. Q J Nucl Med Mol Imaging 2005, 49 (4), 349–60.

8. Liu, L.; Ye, Q.; Wu, Y.; Hsieh, W. Y.; Chen, C. L.; Shen, H. H.; Wang, S. J.; Zhang, H.; Hitchens, T. K.; Ho, C., Tracking T-cells in vivo with a new nano-sized MRI contrast agent. Nanomedicine 2012, 8 (8), 1345–54.

9. Boschi, F.; De Sanctis, F.; Ugel, S.; Spinelli, A. E., T-cell tracking using Cerenkov and radioluminescence imaging. J Biophotonics 2018, 11 (10).

10. Gleich, B.; Weizenecker, J., Tomographic imaging using the nonlinear response of magnetic particles. Nature 2005, 435 (7046), 1214–7.

11. Zhou, X. Y.; Tay, Z. W.; Chandrasekharan, P.; Yu, E. Y.; Hensley, D. W.; Orendorff, R.; Jeffris, K. E.; Mai, D.; Zheng, B.; Goodwill, P. W.; Conolly, S. M., Magnetic particle imaging for radiation-free, sensitive and high-contrast vascular imaging and cell tracking. Curr Opin Chem Biol 2018, 45, 131–138.

12. Graeser, M.; Thieben, F.; Szwargulski, P.; Werner, F.; Gdaniec, N.; Boberg, M.; Griese, F.; Moddel, M.; Ludewig, P.; van de Ven, D.; Weber, O. M.; Woywode, O.; Gleich, B.; Knopp, T., Human-sized magnetic particle imaging for brain applications. Nat Commun 2019, 10 (1), 1936.

13. Savliwala, S.; Chiu-Lam, A.; Unni, M.; Rivera-Rodriguez, A.; Fuller, E.; Sen, K.; Threadcraft, M.; Rinaldi, C., Magnetic nanoparticles. In Nanoparticles for Biomedical Applications: Fundamental Concepts, Biological Interactions and Clinical Applications, Chung, E. J.; Leon, L.; Rinaldi, C., Eds. Elsevier: ProQuest Ebook Central, 2019; pp 195–221.

14. Pouliquen, D.; Le Jeune, J. J.; Perdrisot, R.; Ermias, A.; Jallet, P., Iron oxide nanoparticles for use as an MRI contrast agent: Pharmacokinetics and metabolism. Magnetic Resonance Imaging 1991, 9 (3), 275–283.

15. Weissleder, R.; Stark, D. D.; Engelstad, B. L.; Bacon, B. R.; Compton, C. C.; White, D. L.; Jacobs, P.; Lewis, J., Superparamagnetic iron oxide: pharmacokinetics and toxicity. AJR Am J Roentgenol 1989, 152 (1), 167–73.

16. Cooley, C. Z.; Mandeville, J. B.; Mason, E. E.; Mandeville, E. T.; Wald, L. L., Rodent Cerebral Blood Volume (CBV) changes during hypercapnia observed using Magnetic Particle Imaging (MPI) detection. Neuroimage 2018, 178, 713–720.

17. Kaul, M. G.; Mummert, T.; Jung, C.; Salamon, J.; Khandhar, A. P.; Ferguson, R. M.; Kemp, S. J.; Ittrich, H.; Krishnan, K. M.; Adam, G.; Knopp, T., In vitro and in vivo comparison of a tailored magnetic particle imaging blood pool tracer with Resovist. Phys Med Biol 2017, 62 (9), 3454–3469.

18. Kaul, M. G.; Salamon, J.; Knopp, T.; Ittrich, H.; Adam, G.; Weller, H.; Jung, C., Magnetic particle imaging for in vivo blood flow velocity measurements in mice. Phys Med Biol 2018, 63 (6), 064001.

19. Keselman, P.; Yu, E. Y.; Zhou, X. Y.; Goodwill, P. W.; Chandrasekharan, P.; Ferguson, R. M.; Khandhar, A. P.; Kemp, S. J.; Krishnan, K. M.; Zheng, B.; Conolly, S. M., Tracking short-term biodistribution and long-term clearance of SPIO tracers in magnetic particle imaging. Phys Med Biol 2017, 62 (9), 3440–3453.

20. Ludewig, P.; Gdaniec, N.; Sedlacik, J.; Forkert, N. D.; Szwargulski, P.; Graeser, M.; Adam, G.; Kaul, M. G.; Krishnan, K. M.; Ferguson, R. M.; Khandhar, A. P.; Walczak, P.; Fiehler, J.; Thomalla, G.; Gerloff, C.; Knopp, T.; Magnus, T., Magnetic Particle Imaging for Real-Time Perfusion Imaging in Acute Stroke. ACS Nano 2017, 11 (10), 10480–10488.

21. Zhou, X. Y.; Jeffris, K. E.; Yu, E. Y.; Zheng, B.; Goodwill, P. W.; Nahid, P.; Conolly, S. M., First in vivo magnetic particle imaging of lung perfusion in rats. Phys Med Biol 2017, 62 (9), 3510–3522.

22. Tay, Z. W.; Chandrasekharan, P.; Zhou, X. Y.; Yu, E.; Zheng, B.; Conolly, S., In vivo tracking and quantification of inhaled aerosol using magnetic particle imaging towards inhaled therapeutic monitoring. Theranostics 2018, 8 (13), 3676–3687.

23. Wegner, F.; Buzug, T. M.; Barkhausen, J., Take a Deep Breath - Monitoring of Inhaled Nanoparticles with Magnetic Particle Imaging. Theranostics 2018, 8 (13), 3691–3692.

24. Yu, E. Y.; Chandrasekharan, P.; Berzon, R.; Tay, Z. W.; Zhou, X. Y.; Khandhar, A. P.; Ferguson, R. M.; Kemp, S. J.; Zheng, B.; Goodwill, P. W.; Wendland, M. F.; Krishnan, K. M.; Behr, S.; Carter, J.; Conolly, S. M., Magnetic Particle Imaging for Highly Sensitive, Quantitative, and Safe in Vivo Gut Bleed Detection in a Murine Model. ACS Nano 2017, 11 (12), 12067–12076.

25. Orendorff, R.; Peck, A. J.; Zheng, B.; Shirazi, S. N.; Matthew Ferguson, R.; Khandhar, A. P.; Kemp, S. J.; Goodwill, P.; Krishnan, K. M.; Brooks, G. A.; Kaufer, D.; Conolly, S., First in vivo traumatic brain injury imaging via magnetic particle imaging. Phys Med Biol 2017, 62 (9), 3501–3509.

26. Yu, E. Y.; Bishop, M.; Zheng, B.; Ferguson, R. M.; Khandhar, A. P.; Kemp, S. J.; Krishnan, K. M.; Goodwill, P. W.; Conolly, S. M., Magnetic Particle Imaging: A Novel in Vivo Imaging Platform for Cancer Detection. Nano Lett 2017, 17 (3), 1648–1654.

27. Arami, H.; Teeman, E.; Troksa, A.; Bradshaw, H.; Saatchi, K.; Tomitaka, A.; Gambhir, S. S.; Hafeli, U. O.; Liggitt, D.; Krishnan, K. M., Tomographic magnetic particle imaging of cancer targeted nanoparticles. Nanoscale 2017, 9 (47), 18723–18730.

28. Zheng, B.; von See, M. P.; Yu, E.; Gunel, B.; Lu, K.; Vazin, T.; Schaffer, D. V.; Goodwill, P. W.; Conolly, S. M., Quantitative Magnetic Particle Imaging Monitors the Transplantation, Biodistribution, and Clearance of Stem Cells In Vivo. Theranostics 2016, 6 (3), 291–301.

29. Zheng, B.; Vazin, T.; Goodwill, P. W.; Conway, A.; Verma, A.; Saritas, E. U.; Schaffer, D.; Conolly, S. M., Magnetic Particle Imaging tracks the long-term fate of in vivo neural cell implants with high image contrast. Sci Rep 2015, 5, 14055.

30. Reilly, K. M.; Loisel, D. A.; Bronson, R. T.; McLaughlin, M. E.; Jacks, T., Nf1;Trp53 mutant mice develop glioblastoma with evidence of strain-specific effects. Nat Genet 2000, 26 (1), 109–13.

31. Pohl-Guimaraes, F.; Yang, C.; Dyson, K. A.; Wildes, T. J.; Drake, J.; Huang, J.; Flores, C.; Sayour, E. J.; Mitchell, D. A., RNA-Modified T Cells Mediate Effective Delivery of Immunomodulatory Cytokines to Brain Tumors. Mol Ther 2019, 27 (4), 837–849.

32. Fedorov, A.; Beichel, R.; Kalpathy-Cramer, J.; Finet, J.; Fillion-Robin, J. C.; Pujol, S.; Bauer, C.; Jennings, D.; Fennessy, F.; Sonka, M.; Buatti, J.; Aylward, S.; Miller, J. V.; Pieper, S.; Kikinis, R., 3D Slicer as an image computing platform for the Quantitative Imaging Network. Magn Reson Imaging 2012, 30 (9), 1323–1341.

33. Johnson, L. D.; Jameson, S. C., Self-specific CD8+ T cells maintain a semi-naive state following lymphopenia-induced proliferation. J Immunol 2010, 184 (10), 5604–11.

34. Prins, R. M.; Shu, C. J.; Radu, C. G.; Vo, D. D.; Khan-Farooqi, H.; Soto, H.; Yang, M. Y.; Lin, M. S.; Shelly, S.; Witte, O. N.; Ribas, A.; Liau, L. M., Anti-tumor activity and trafficking of self, tumor-specific T cells against tumors located in the brain. Cancer Immunol Immunother 2008, 57 (9), 1279–89.

35. Flores, C.; Pham, C.; Snyder, D.; Yang, S.; Sanchez-Perez, L.; Sayour, E.; Cui, X.; Kemeny, H.; Friedman, H.; Bigner, D. D.; Sampson, J.; Mitchell, D. A., Novel role of hematopoietic stem cells in immunologic rejection of malignant gliomas. Oncoimmunology 2015, 4 (3), e994374.

36. Reimer, P.; Balzer, T., Ferucarbotran (Resovist): a new clinically approved RES-specific contrast agent for contrast-enhanced MRI of the liver: properties, clinical development, and applications. Eur Radiol 2003, 13 (6), 1266–76.

37. Yang, C. Y.; Tai, M. F.; Lin, C. P.; Lu, C. W.; Wang, J. L.; Hsiao, J. K.; Liu, H. M., Mechanism of cellular uptake and impact of ferucarbotran on macrophage physiology. Plos One 2011, 6 (9), e25524.

38. Namkung, S.; Zech, C. J.; Helmberger, T.; Reiser, M. F.; Schoenberg, S. O., Superparamagnetic iron oxide (SPIO)-enhanced liver MRI with ferucarbotran: efficacy for characterization of focal liver lesions. J Magn Reson Imaging 2007, 25 (4), 755–65.

39. Baars, P. A.; Sierro, S.; Arens, R.; Tesselaar, K.; Hooibrink, B.; Klenerman, P.; van Lier, R. A., Properties of murine (CD8+)CD27-T cells. Eur J Immunol 2005, 35 (11), 3131–41.

40. Suzuka, H.; Mimura, A.; Inaoka, Y.; Murase, K., Magnetic Nanoparticles in Macrophages and Cancer Cells Exhibit Different Signal Behavior on Magnetic Particle Imaging. J Nanosci Nanotechnol 2019, 19 (11), 6857–6865.

41. Chandrasekharan, P.; Tay, Z. W.; Zhou, X. Y.; Yu, E.; Orendorff, R.; Hensley, D.; Huynh, Q.; Fung, K. L. B.; VanHook, C. C.; Goodwill, P.; Zheng, B.; Conolly, S., A perspective on a rapid and radiation-free tracer imaging modality, magnetic particle imaging, with promise for clinical translation. Br J Radiol 2018, 91 (1091), 20180326.

42. Chapelin, F.; Capitini, C. M.; Ahrens, E. T., Fluorine-19 MRI for detection and quantification of immune cell therapy for cancer. J Immunother Cancer 2018, 6 (1), 105.

43. Gonzales, C.; Yoshihara, H. A.; Dilek, N.; Leignadier, J.; Irving, M.; Mieville, P.; Helm, L.; Michielin, O.; Schwitter, J., In-Vivo Detection and Tracking of T Cells in Various Organs in a Melanoma Tumor Model by 19F-Fluorine MRS/MRI. Plos One 2016, 11 (10), e0164557.

44. Visioni, A.; Kim, M.; Wilfong, C.; Blum, A.; Powers, C.; Fisher, D.; Gabriel, E.; Skitzki, J., Intra-arterial Versus Intravenous Adoptive Cell Therapy in a Mouse Tumor Model. J Immunother 2018, 41 (7), 313–318.

45. Armbruster, D. A.; Pry, T., Limit of blank, limit of detection and limit of quantitation. Clin Biochem Rev 2008, 29 Suppl 1, S49–52.

