## Supplementary Information for "Tracking Adoptive T Cell Therapy Using Magnetic Particle Imaging"

**SUPPORTING INFORMATION**

**Supplemental methodology:**

**MPI Limit of Detection:** To determine the limit of detection (LoD) in our MPI system, a dilution series of ferucarbotran tracer was prepared, with decreasing iron mass by a factor of 2. All scanned samples consisted of a volume of 1 µL in a 0.2 mL microcentrifuge tube. Samples (n=3) were imaged at two different positions within the MPI field of view. MPI scans were performed using the Momentum™ imager from Magnetic Insight (Alameda, CA, USA) in the high-sensitivity (3 T/m) scan mode. The images were analyzed using VivoQuant and Matlab to evaluate signal intensity in regions of interest containing the tracer samples. The LoD was calculated by analyzing the background signal from empty scans, the limit of blank (LoB), and the maximum intensity signal of samples at low concentrations.[45]

$$LoB = mean_{blank} + 1.645(SD_{blank})$$

$$LoD = LoB + 1.645(SD_{low\ concentration\ sample})$$

**MATLAB image processing:** DICOM files from the Momentum™ imager were stored with a linear transformation. In order to visually compare images, a MATLAB code was written to rescale images to their original representation, and to remove background noise. Background noise was defined as the signal intensity detected in the region where no sample was present.

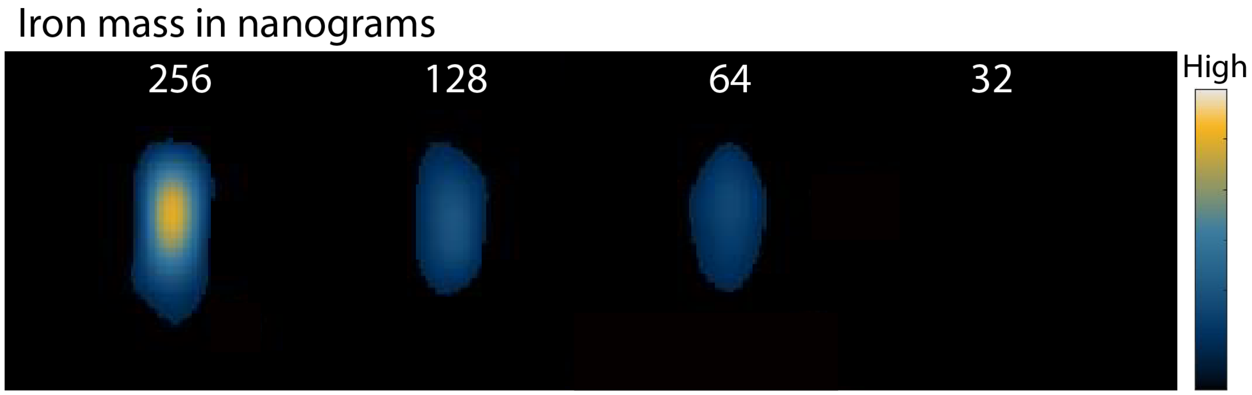


**Figure S1.** MPI signal intensity of free ferucarbotran tracer. A dilution series of known iron mass shows that signal can be detected at 64 ng with a calculated limit of detection of 38 ng of iron.

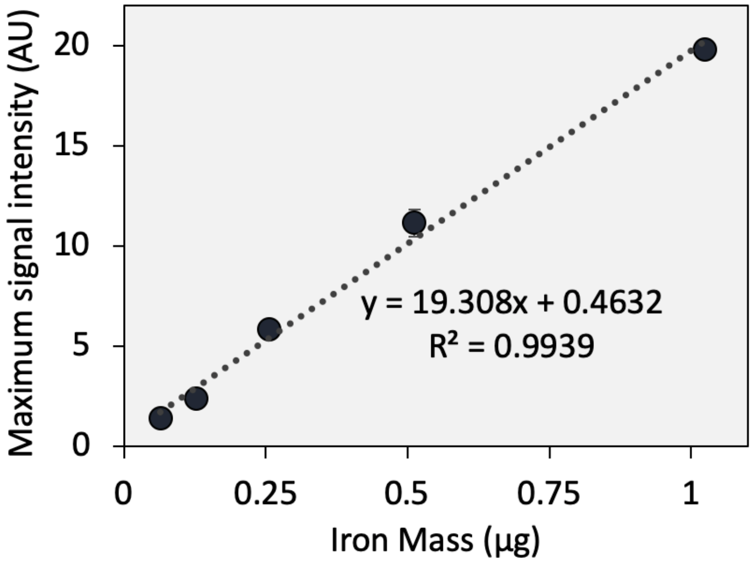


**Figure S2.** Linear correlation of ferucarbotran iron mass and MPI signal intensity. Note: Error bars are smaller than the markers.

**Table S1.** Values used to determine the limit of detection (LoD) for ferucarbotran.

| | MPI signal |
|---|---|
| $Mean_{Blank}$ | -0.013 |
| $SD_{Blank}$ | 0.269 |
| LoB | 0.429 |
| $SD_{Max\ Signal\ at\ 64\ ngFe}$ | 0.108 |
| LoD | 0.608 |

For the dilution series of ferucarbotran, the signal intensity decreases with decreasing iron mass (**Figure S1** and **Figure S2)**. The high-sensitivity scan mode used throughout this paper has a low gradient field strength of 3 T/m. Low gradient fields do not fully saturate the particles in the vicinity of the field free region, and more signal can be obtained at the expense of resolution. The calculated limit of detection is approximately 38 ng of iron for ferucarbotran nanoparticles, which is consistent with the lowest number of ferucarbotran-labeled T cells (~50,000 cells) that were detect *in vitro*. The LoD in terms of iron mass was calculated using the LoD in signal intensity from **Table S1**, and the value was correlated to iron mass using the regression line from **Figure S2**.

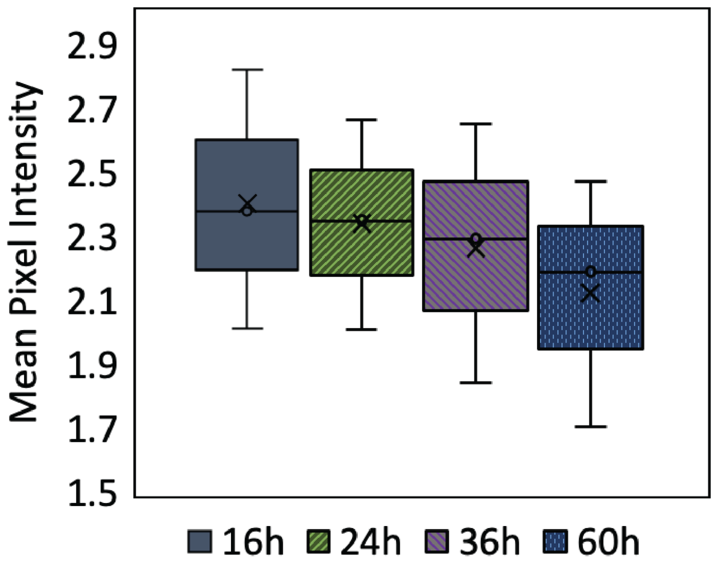


**Figure S3.** Mean pixel intensity values for intraventricular injection of ferucarbotran-labeled T cells in mouse (n=3).

After intraventricular injection of ferucarbotran-labeled T cells and longitudinal *in vivo* MPI, regions of interest were drawn around the MPI signals in each mouse (n=3) and mean pixel intensity was obtained. **Figure 3S** shows a box plot of the mean pixel intensity values. A small decrease in signal is observed over time. However, at the studied time points, these changes are not statically significant at a p-value of 0.01.
